# Rare alternative second line injectable drug resistance markers identified by gene-wise genome wide association in *M. tuberculosis* with unexplained resistance

**DOI:** 10.1101/2021.11.23.469801

**Authors:** Derek Conkle-Gutierrez, Calvin Kim, Sarah M. Ramirez-Busby, Samuel J. Modlin, Mikael Mansjö, Jim Werngren, Leen Rigouts, Sven E. Hoffner, Faramarz Valafar

## Abstract

Point mutations in the *rrs* gene and *eis* promoter are known to confer resistance to second-line injectable drugs (SLIDs) amikacin (AMK), capreomycin (CAP), and kanamycin (KAN). While mutations in these canonical genes confer a majority of SLID-resistance, alternative mechanisms of resistance are not uncommon and threaten effective treatment decisions when using conventional molecular diagnostics. In total, 1184 clinical *M. tuberculosis* isolates from 7 countries were studied for genomic markers associated with phenotypic resistance. The markers *rrs:*A1401G and *rrs:*G1484T were associated with resistance to all three SLIDs, and three known markers in the *eis* promoter (*eis*:G-10A, *eis*:C-12T, and *eis*:C-14T) were similarly associated with kanamycin resistance (KAN-R). Among 325, 324, 270 AMK-R, CAP-R, and KAN-R isolates, 264 (81.2%), 250 (77.2%), and 249 (92.3%) harbored canonical mutations, respectively. Thirteen isolates harbored more than one canonical mutation. Canonical mutations did not account for 111 of the phenotypically resistant isolates. A gene-wise method identified three genes and promoters with mutations that, on aggregate, associated with unexplained resistance to at least one SLID. Our analysis associated *whiB7* promoter mutations with KAN resistance, supporting clinical relevance for the previously demonstrated role of *whiB7* overexpression in KAN resistance. We also provide evidence for the novel association of *ppe51* (a gene previously associated with various antimicrobial compounds) with AMK resistance, and for the novel association of *thrB* with AMK and CAP resistance. The use of gene-wise association can provide additional insight, and therefore is recommended for identification of rare mechanisms of resistance when individual mutations carry insufficient statistical power.

## Introduction

Tuberculosis (TB) remains a constant global public health threat due to rising cases of drug resistance among various strains of *Mycobacterium tuberculosis*. Half a million estimated TB cases were rifampicin resistant in 2020, including 3-4% of new TB cases and 18-21% of previously treated cases^1^. This trend is exacerbated in countries of the former Soviet Union, where over half of previously treated TB patients were rifampicin resistant^1^. In 2018, 78% of rifampicin resistant cases were also resistant to isoniazid, making them multidrug resistant tuberculosis (MDR-TB)^2^. In 2018, an estimated 6.2% of MDR-TB cases were extensively drug resistant (XDR), then defined as MDR-TB strains that were additionally resistant to at least a fluoroquinolone and a second-line injectable drug^2^ (SLID).

Successful Tuberculosis treatment relies on early identification and effective regimens, which can be ensured by rapid and accurate drug-susceptibility testing (DST) to identify potential MDR-TB cases. MDR-TB should be treated with combinations of drugs shown *in vitro* to be effective^3^. Phenotypic DST takes weeks, during which time patients may face ineffective treatment regimens with often debilitating side effects. Molecular diagnostics in contrast can be performed rapidly. As these rely on genetic markers of resistance, to improve their accuracy we must understand the mechanisms behind the resistance and comprehensively identify all their markers.

The SLIDs amikacin (AMK) and kanamycin (KAN) kill *M. tuberculosis* cells by binding to the 16S ribosomal subunit and by disrupting translation^4^. The bactericidal mechanism of the SLID capreomycin (CAP) is less understood. It is hypothesized to bind to the 70S ribosomal subunit and limit mRNA-tRNA translocation^5^. In 2018, the WHO recommended against the use of CAP or KAN due to their side effects, such as ototoxicity and nephrotoxicity^6^, and their significant association with treatment failure^7^. Even though AMK is now being phased out in favor of bedaquiline^8^, it is still widely used. Although CAP and KAN are currently not recommended in treatment regimens, understanding mechanisms of resistance to these drugs can inform and corroborate our understanding of AMK-resistance (AMK-R), as the three drugs have similar mechanisms of action.

Genetic markers can rapidly identify SLID-resistance (SLID-R) in clinical isolates^9^. Across multiple studies, the most frequently observed SLID-R marker within the *M. tuberculosis* complex is *rrs*:A1401G^10–13^. This single nucleotide polymorphism (SNP) in the 16S ribosomal RNA (rRNA) gene *rrs* typically causes cross-resistance to all three SLIDs^11,13^. Other mutations in *rrs* also associate with SLID-R, but are far more rare^14^.

While *rrs*:A1401G typically causes resistance to all three SLIDs, not all isolates resistant to one or more SLIDs have this marker^11^. Many KAN-resistant (KAN-R) isolates, for example, harbor variants in the promoter of *eis*^11,15,16^. The *eis* gene encodes an N-acetyltransferase, which can inactivate KAN when overexpressed^15^. Spectrophotometric assays have further shown that *eis* acetylates KAN^17^. These *eis* promoter markers are common in *M. tuberculosis* strains from the former Soviet Union, where the use of KAN has been high^16^. Together with *rrs*:A1401G and *rrs*:G1484T, these known markers (Table 1) explain most SLID-R *M. tuberculosis* isolates^18^.

**Table 1.**
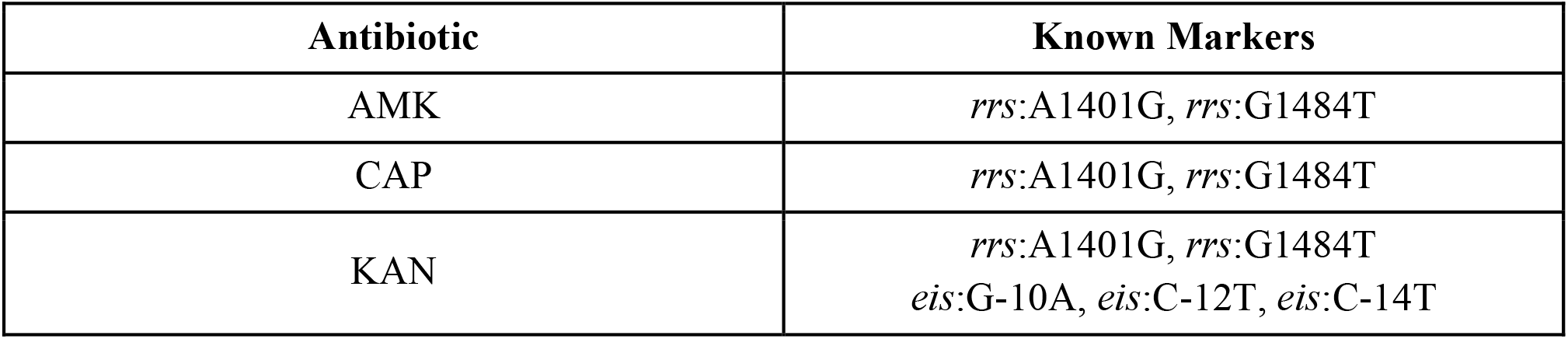
The known SLID-R markers in *M. tuberculosis*. This set of mutations was derived from a previous study^18^, and used to determine the expected resistance of clinical isolates to AMK, CAP, and KAN.

Previous studies have found SLID-R isolates with no known markers^11,13^. Mutations in several genes have been suggested to cause resistance in these isolates, including *tlyA*^19^, *whiB7*^20^, *vapC21*^21^, and *bfrB*, also known as *ferritin*^22,23^. Knockout variants in *tlyA* induce CAP resistance (CAP-R) *in vitro*^5^. Loss-of-function mutations in *tlyA*^5,18,19^ are proposed to prevent CAP from binding to the 16S subunit^19^ by preventing methylation of the 16S^24^.

To find alternative mechanisms of SLID-R, we performed a genome wide association study (GWAS) on 1184 clinical *M. tuberculosis* isolates, including 111 SLID-R isolates with no known SLID-R markers. Our methods corroborated the association of *whiB7* with kanamycin resistance, and identified several putative amikacin resistance markers in *ppe51*, a transport mediator previously implicated in resistance to drug candidate 3,3-bis-di(methylsulfonyl)propionamide^25^.

## Results

This study surveyed 1184 clinical isolates for injectable resistance markers. AMK and CAP phenotypic DST data were available for 1163 and 1159 isolates, respectively (Table 2). Only 496 isolates had phenotypic DST data available for KAN (Table 2). Our set of clinical isolates included 333 isolates sequenced on Single Molecule Real Time (SMRT) sequencers, which can resolve many known blind spots in the *M. tuberculosis* genome, such as in the PE and PPE gene families^26^. The isolates were categorized based on phenotypic DST and the presence of known resistance markers.

**Table 2.**
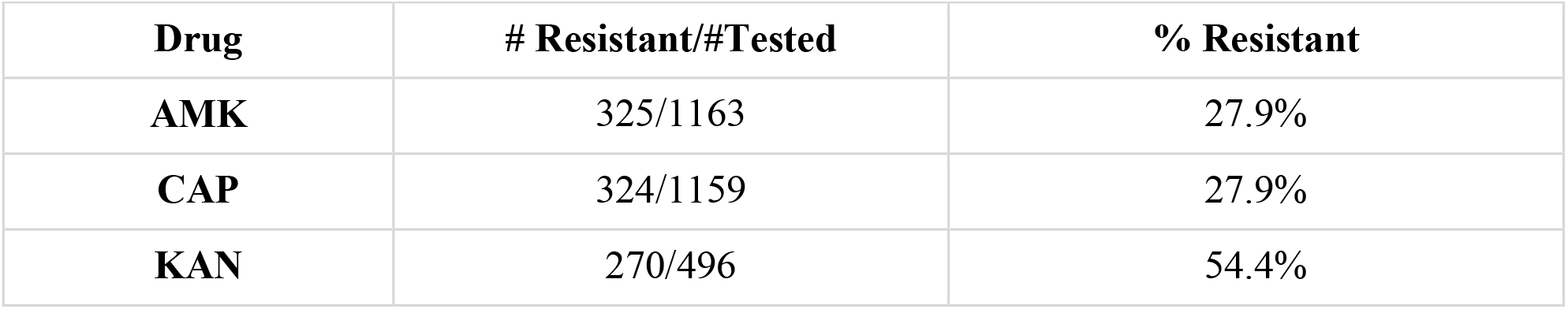
Phenotypic Drug Susceptibility Testing Results. Number of total resistant *M. tuberculosis* isolates out of all isolates with available phenotypic drug susceptibility testing (DST) for each drug, and the percentage of isolates with DST to a given drug that show resistance.

### Mutations in rrs and eis Promoter Associate with SLID Resistance and Specific Lineages

Known SLID-R markers in the *rrs* gene and *eis* promoter associated strongly with resistance (Table 3). They predicted SLID-R with similar sensitivity to a 2018 GWAS study^27^, though the sensitivity was lower than in earlier studies^5,18^. The lower sensitivity is likely due to the deliberate selection of clinical isolates with discordant genotypes and phenotypes. Variant *rrs:*A1401G was the most frequent marker (N = 260, Table 4), while *rrs:*G1484T was the least frequent (N = 4, Table 4). However, no isolate with *rrs:*G1484T was susceptible to any SLID they were tested for (Table 4). The known marker *rrs:*A1401G was more frequent (181/700) within Lineage 2 (East-Asian) than within any other lineage (Figure 1, Table S1).

**Table 3.** Sensitivity and specificity of SLID-R prediction with known markers. A 95% confidence interval for each estimate was calculated using the score method with continuity correction^28^. Isolates with genotypic-phenotypic concordance were classified as “explained”. Otherwise, they are “unexplained”.

| Drug | Explained Resistant | Explained Susceptible | Unexplained Resistant | Unexplained Susceptible | Sensitivity (% [95% CI]) | Specificity (% [95% CI]) |
| --- | --- | --- | --- | --- | --- | --- |
| AMK | 264 | 829 | 61 | 9 | 81.2<br>(78.8 - 83.4) | 98.9<br>(98.1 - 99.4) |
| CAP | 250 | 813 | 74 | 22 | 77.2<br>(74.6 - 79.5) | 97.4<br>(96.2 - 98.2) |
| KAN | 249 | 223 | 21 | 3 | 92.3<br>(89.6 - 94.4) | 98.7<br>(97.1 - 99.4) |

**Table 4.** Number of *M. tuberculosis* isolates with each known marker. Isolates are classified based on their resistance or susceptibility to each SLID. R=resistant; S=sensitive. Note that some isolates carried multiple markers.

| Variant | AMK-R | AMK-S | CAP-R | CAP-S | KAN-R | KAN-S |
| --- | --- | --- | --- | --- | --- | --- |
| *rrs*:A1401G | 260 | 9 | 246 | 22 | 188 | 1 |
| *rrs*:G1484T | 4 | 0 | 4 | 0 | 3 | 0 |
| *eis*:G-10A | 5 | 49 | 3 | 51 | 10 | 0 |
| *eis*:C-12T | 10 | 95 | 19 | 84 | 42 | 2 |
| *eis*:C-14T | 11 | 18 | 3 | 26 | 10 | 0 |

**Figure 1.**
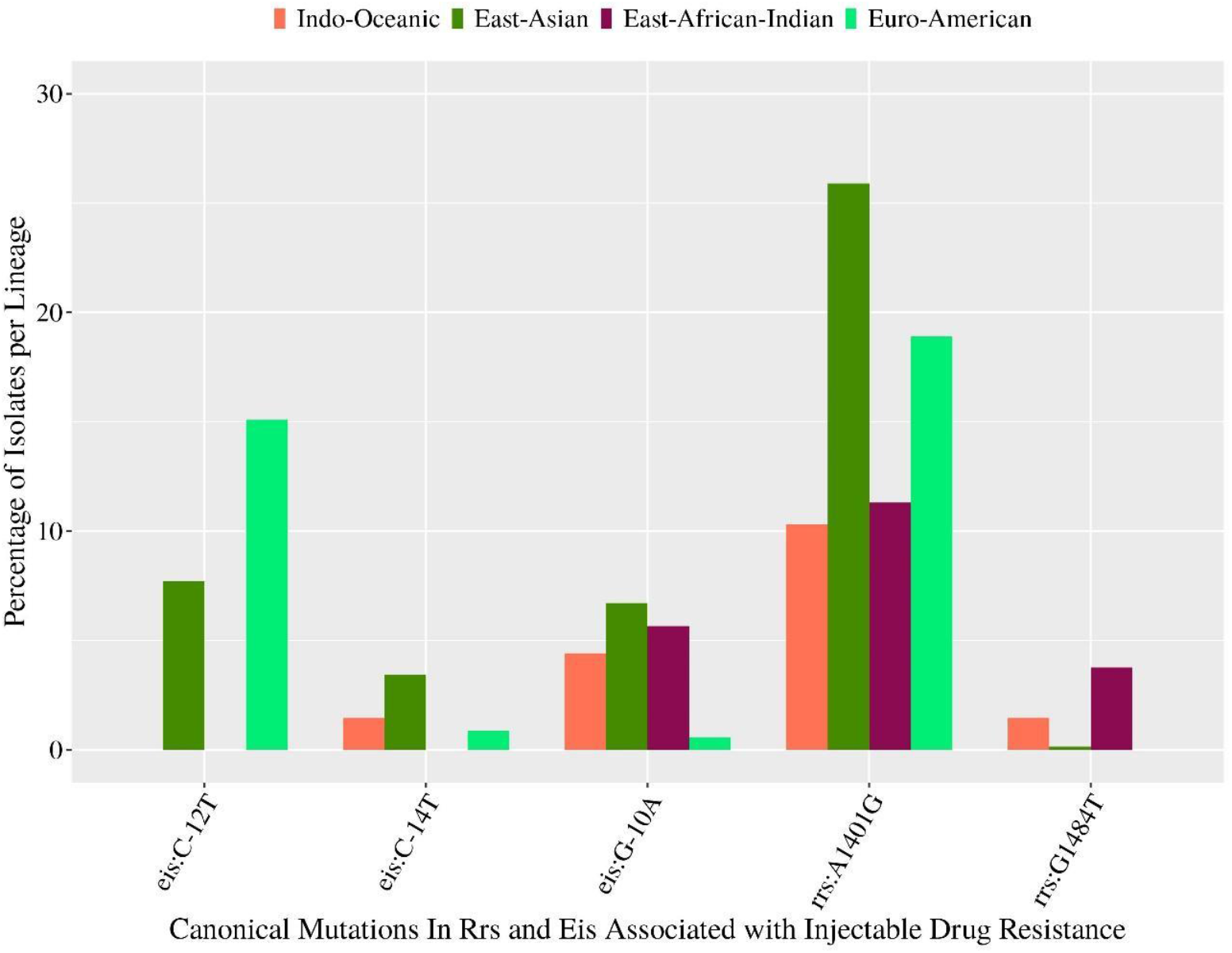
Frequency of known resistance markers within each lineage among 1184 clinical *M. tuberculosis* isolates. The Y-axis represents the percent of clinical *M. tuberculosis* isolates from each lineage that possess the mutations represented on the X-axis. The lineage of 12 isolates could not be determined, including one isolate with *eis*:C-14T, and one isolate with *rrs*:A1401G. One other isolate belonged to the *M. africanum* species, West African 1 lineage.

Ten unexplained AMK-R isolates could be explained by the *eis* promoter mutation C-14T. The mutation was carried by 11 AMK-R and 18 AMK-S isolates (Table 3). Ten of the AMK-R isolates with *eis*:C-14T carried no SLID-R marker in *rrs* (Table 3, S2). Another three unexplained AMK-R isolates had evidence of heteroresistance, with *rrs*:A1401G supported by 10-20% of the mapped reads (Table S3). No unexplained AMK-R isolates had reads supporting *rrs*:G1484T.

Mutations in the *eis* promoter were especially common in isolates from Moldova (Table 5). Among the 270 KAN-R isolates, isolates with known KAN-R markers in the *eis* promoter were 27.8 times more likely to be from Moldova than isolates without known markers in the *eis* promoter (95% CI 12.8-64.8, p-value < 2.2e-16, Fisher’s Exact Test, Table 5). The 89 isolates from Moldova belonged exclusively to Lineage 4 (Euro-American, n = 51) and East-Asian (n = 38). Similarly, among all 1184 isolates, the only isolates with variant *eis*:C-12T were East-Asian (54/700) or Euro-American (53/350, Figure 1). Two isolates carried both *eis*:C-12T and another known marker in the *eis* promoter (Table S2). Ten isolates with known markers in the *eis* promoter also carried the marker *rrs*:A1401G (Table S2).

**Table 5.** Geographic specificity of KAN Resistance markers in *eis* promoter. Contingency tables reporting the association between possession of a known KAN-R marker in the *eis* promoter and collection from Moldova, stratified by KAN DST. Two Moldovan isolates had low sequencing coverage in the *eis* promoter, and were thus excluded from this contingency table, and all KAN analysis. The number of isolates with either of the SLID-R markers *rrs*:A1401G and *rrs*:G1484T are included in parenthesis.

| Number of Kanamycin-Resistant Isolates |  |  |  |
| --- | --- | --- | --- |
|  | Moldovan | Not Moldovan | Totals |
| Carries Known Marker in *eis* Promoter | 48 (2) | 13 (1) | 61 (3) |
| No Known Marker in *eis* Promoter | 24 (21) | 185 (170) | 209 (191) |
| Totals | 72 | 198 | 270 |
| Number of Kanamycin-Susceptible Isolates |  |  |  |
|  | Moldovan | Not Moldovan | Totals |
| Carries Known Marker in *eis* Promoter | 2 | 0 | 2 |
| No Known Marker in *eis* Promoter | 13 | 211 | 224 |
| Totals | 15 | 211 | 226 |

### Mutations rrs:A1401G and rrs:G1484T Associate with Cross-Resistance

Phenotypic DST results were available for all three SLIDs in 476 isolates, of which 193 isolates were resistant to all three drugs (Figure 2). Known marker *rrs:*A1401G explained 181 of the 193 cross-resistant isolates (Table 6). Of the remaining twelve cross-resistant isolates without *rrs:*A1401G, three isolates had known marker *rrs:*G1484T (Table 6). No isolate had both *rrs:*A1401G and *rrs:*G1484T (Table S2).

**Table 6.**
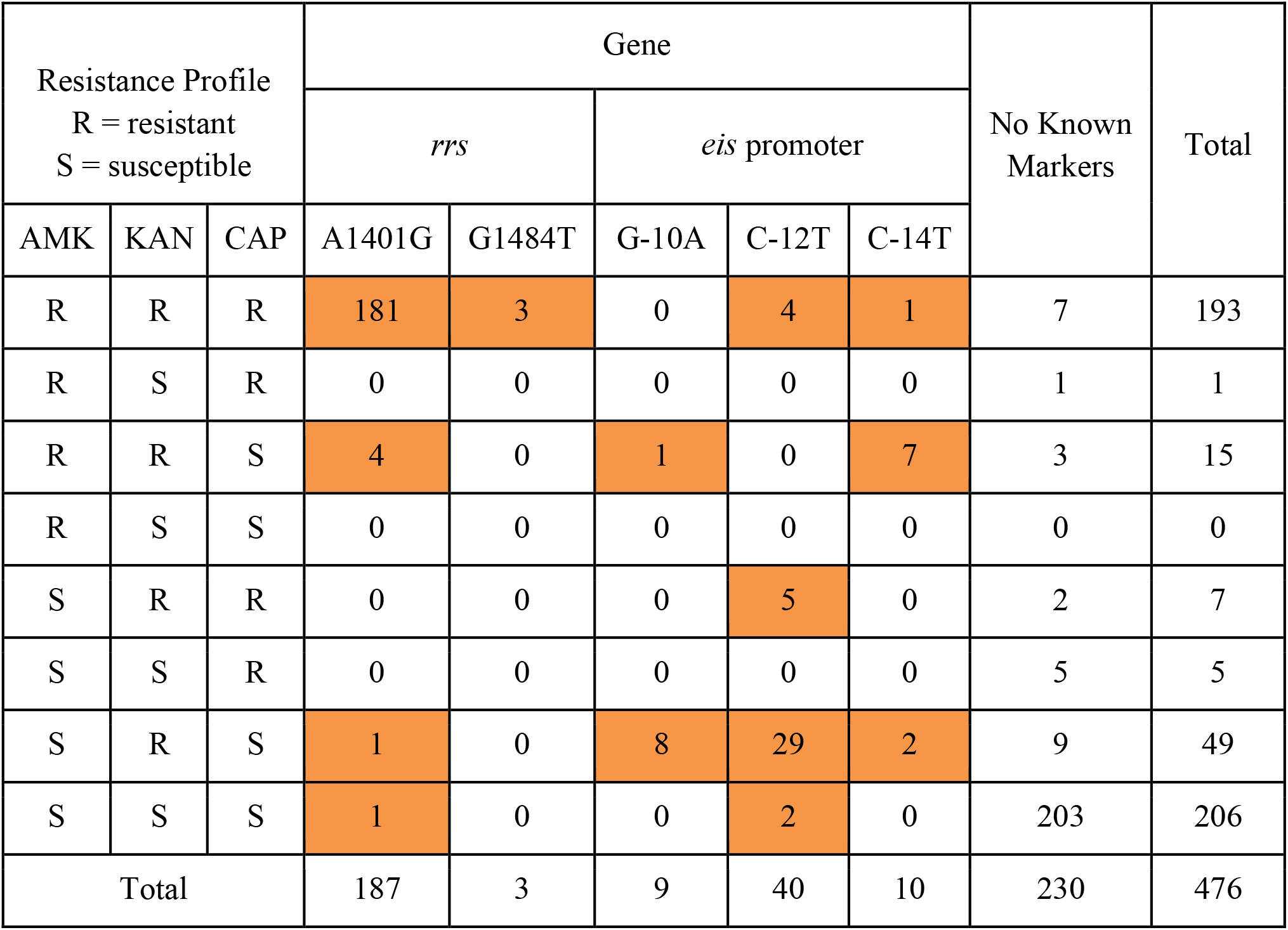
Association of SLID-R profiles to known resistance markers. Each row represents one of the eight possible resistance profiles to the three SLIDs. The columns under *rrs* and *eis* promoter report the number of isolates carrying each known SLID-R marker with each resistance profile. As the matrix is sparse, cells with nonzero values are highlighted. The “No Known Markers” column reports the number of isolates carrying no known SLID-R marker, with each drug resistance profile. Only the 476 isolates with phenotypic DST results for all three SLIDs were included in these counts. Note that 3 of these isolates carried multiple known markers, all of them fully SLID cross-resistant.

| Resistance Profile<br>R = resistant<br>S = susceptible |  |  | Gene |  |  |  |  | No Known Markers | Total |
| --- | --- | --- | --- | --- | --- | --- | --- | --- | --- |
|  |  |  | *rrs* |  | *eis* promoter |  |  |  |  |
| AMK | KAN | CAP | A1401G | G1484T | G-10A | C-12T | C-14T |  |  |
| R | R | R | 181 | 3 | 0 | 4 | 1 | 7 | 193 |
| R | S | R | 0 | 0 | 0 | 0 | 0 | 1 | 1 |
| R | R | S | 4 | 0 | 1 | 0 | 7 | 3 | 15 |
| R | S | S | 0 | 0 | 0 | 0 | 0 | 0 | 0 |
| S | R | R | 0 | 0 | 0 | 5 | 0 | 2 | 7 |
| S | S | R | 0 | 0 | 0 | 0 | 0 | 5 | 5 |
| S | R | S | 1 | 0 | 8 | 29 | 2 | 9 | 49 |
| S | S | S | 1 | 0 | 0 | 2 | 0 | 203 | 206 |
| Total |  |  | 187 | 3 | 9 | 40 | 10 | 230 | 476 |

**Figure 2.**
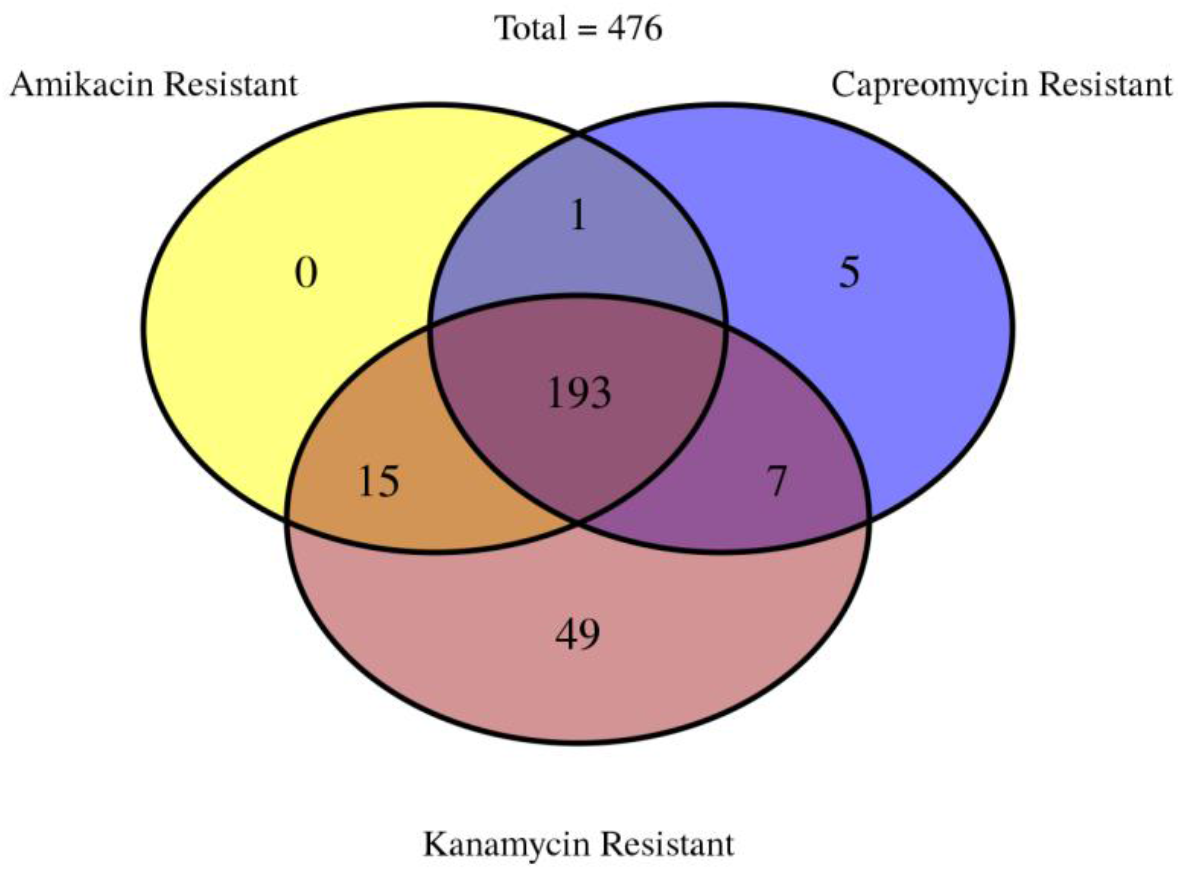
Cross resistance among Second-line Injectable Drugs (SLIDs). Venn Diagram reporting the number of *M. tuberculosis* clinical isolates with overlapping resistance across all three SLIDs. Only the 476 isolates with phenotypic DST results for all three SLIDs were included in this figure.

The known marker *rrs*:A1401G did not always confer full cross-resistance. Four isolates with *rrs:*A1401G were AMK-R and KAN-R, yet CAP-S (Table 6). In total, 22 isolates with *rrs:*A1401G were CAP-S (Table 4). All 22 of these CAP-discordant isolates were AMK-R (KAN DST was not available for 18 of them).

Isolates with full SLID cross-resistance harbored known markers 97.45 (95% CI 43.26-2258.38, p-value < 2.2e-16, Fisher’s Exact Test) times more often than isolates without full cross-resistance (Table 6, Table S4). While no isolate was mono-resistant to AMK, one isolate was simultaneously AMK-R, CAP-R, and KAN-S (Figure 2, Table 6). This abnormal resistance profile may be the result of a mistake in phenotypic DST. The isolate carried a single *rrs* variant, *rrs:*C492T, a mutation present in another eighteen isolates, all susceptible to all three SLIDs (Figure 3). Fifteen isolates were AMK-R, KAN-R and CAP-S, of which four carried *rrs*:A1401G, eight carried markers in the *eis* promoter, and three carried no known markers (Figure 2, Table 6).

**Figure 3.**
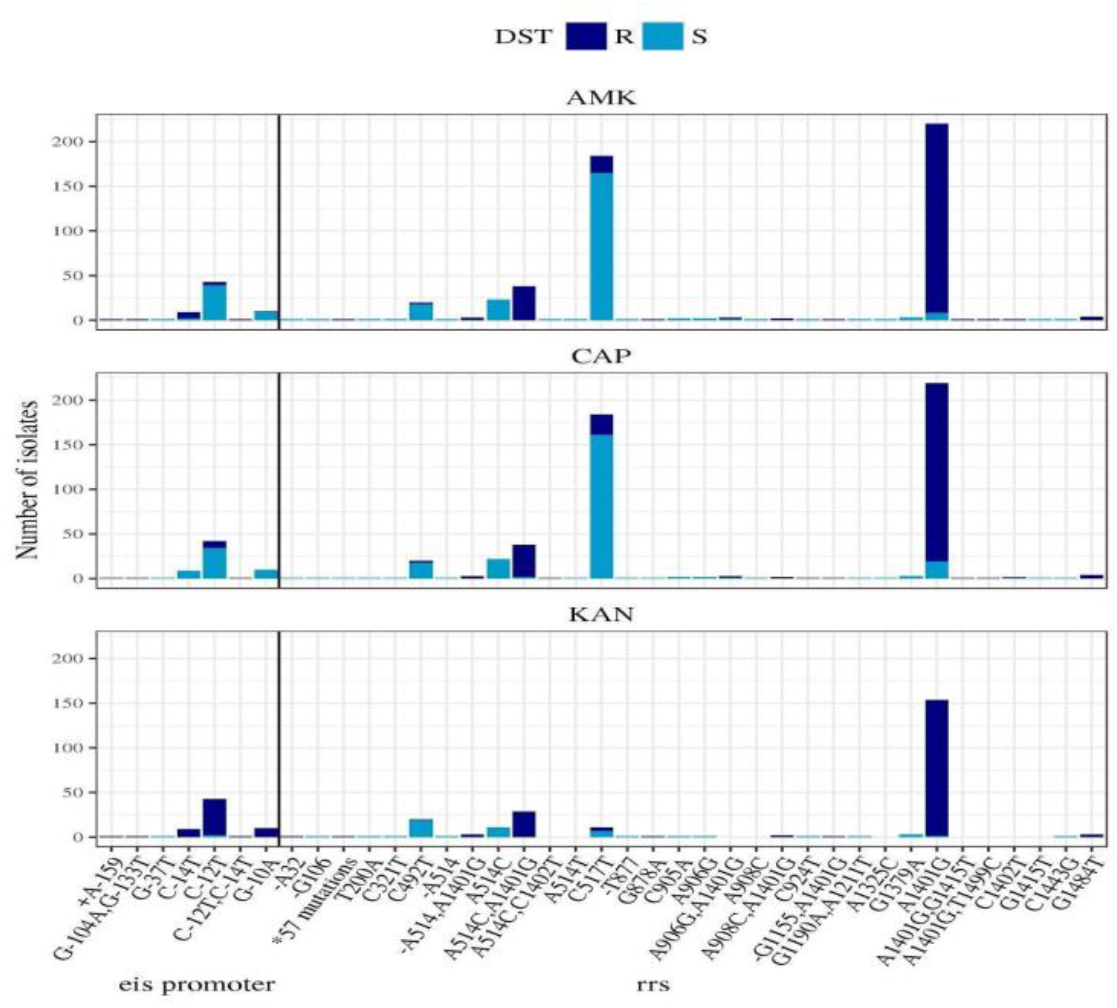
Mutations in *eis* promoter and *rrs*. Number of clinical *M. tuberculosis* isolates with mutations in the *rrs* gene or *eis* promoter, with resistance or susceptibility to AMK, CAP, and KAN. Each column reports the number of resistant (R, dark blue bar) and susceptible (S, light blue bar) isolates carrying each mutation. For each drug, only isolates with phenotypic DST for that drug were counted. A vertical line separates the mutations found in *rrs* from those in the *eis* promoter. Note that column “*57 mutations” represents a set of 57 *rrs* variants called in a single isolate, which were combined for brevity.

### High Co-occurrence of rrs:A1401G with Streptomycin Resistance Marker rrs:A514C

Variants *rrs:*C492T, *rrs:*C517T, and *rrs:*A514C were the most frequent non-canonical *rrs* mutations among the clinical isolates (Figure 3). The three mutations did not associate with resistance to any SLID. Both *rrs*:A514C and *rrs*:C517T are known streptomycin resistance markers^29^. Variant *rrs*:A514C was frequently carried by isolates that also carried the SLID-R marker *rrs:*A1401G. Among all clinical isolates, *rrs:*A514C was 6.0 times more likely to be carried in isolates with *rrs:*A1401G than in isolates without *rrs:*A1401G (p-value = 2.424e-11, Fisher’s Exact Test).

### Insufficient Statistical Power to Identify Individual Alternative Resistance Markers

As there were only 21 unexplained KAN-R isolates in this study (Table 3), there was insufficient statistical power to identify potential rare KAN-R mechanisms. In this isolate set, aside from known markers, no significant association was found between KAN-R and any other mutations in the *eis* promoter. While prior studies found the mutation *eis:*G-37T in isolates with at least low level resistance to KAN^15,30^, only one of our isolates carried it (Figure 3). This isolate was phenotypically KAN-S, though it is possible that MIC would reveal low level resistance to KAN. Similarly, the mutation *rrs*:C1402T was carried by only three isolates, including both 1 AMK-R, 1 AMK-S, 1 AMK untested, 2 CAP-R, and 1 CAP-S isolates (Figure 3).

Mutations in the RNA methylase *tlyA* confer CAP-R *in vitro*^5^ and have previously been observed in clinical isolates^31^. However, in this isolate set no significant association was found between CAP-R and any mutation in *tlyA*. Only six non-synonymous *tlyA* mutations were carried by any CAP-R isolates that did not also carry the known marker *rrs:*A1401G (Figure S1). None of these six mutations were carried by more than two CAP-R isolates (Figure S1).

Excluding isolates with known SLID-R markers, no individual mutation in the genome predicted resistance to any SLID with greater than 1.34% sensitivity in this isolate set (Table 7, Table S5). Several mutations were completely absent from susceptible isolates, though were only carried by at most four resistant isolates (Table 7). These mutations may be alternative SLID-R mechanisms. However due to their rarity, there was insufficient statistical power to determine this through association.

**Table 7.** Genome wide association study, top individual variants. The most frequent variants in ‘unexplained resistant’ isolates, SLID-R isolates that lack known resistance markers. Variants carried by one or more susceptible isolate were excluded. For each mutation, the column “Unexplained Resistant Isolates” counts the number of SLID-R isolates lacking known resistance markers that carry the mutation. The columns “Sensitivity” and “Specificity” report the sensitivity and specificity of predicting drug resistance using only the mutation in the “Mutation” column. Separate sensitivity and specificity values were calculated for each SLID. Only SLID-S and unexplained SLID-R isolates were included.

| Gene | Mutation | SLID | Sensitivity (%) | Specificity (%) | Unexplained Resistant Isolates |
| --- | --- | --- | --- | --- | --- |
| *Rv2815c-Rv2816c* | CT3122937C | AMK | 0.85 | 100 | 3 |
| *thrB_prom* | GAT-97G | AMK | 0.57 | 100 | 2 |
| *pe_pgrs30* | G1682-1683GC | AMK | 0.57 | 100 | 2 |
| *Rv2815c_prom* | G-61GA | CAP | 0.58 | 100 | 2 |
| *ppe23* | F248S | CAP | 0.58 | 100 | 2 |
| *pe_pgrs30* | G1682-1683GC | CAP | 0.58 | 100 | 2 |
| *pe_pgrs57* | ACCG2072-2074A | KAN | 1.34 | 100 | 4 |
| *Rv0988* | L191A | KAN | 1.34 | 100 | 4 |
| *pe_pgrs54* | D1579T | KAN | 1.34 | 100 | 4 |

Several genes have previously been suggested to affect SLID-R. Support vector machine approach identified variants in three genes as potential determinants of XDR phenotype: *vapC21, Rv3471c*, and *Rv3848*^21^. Comparative proteomics suggested 12 genes may be involved in resistance to AMK or KAN: *atpA, tig, lpdC, tuf, moxR1, Rv2005c, 35kd_ag, prcA, Rv0148, bfrA, bfrB, hspX*^22^. Efflux pump *Rv1258c*^32^, transcriptional regulator *whiB7*^33^, and virulence gene *whiB6*^34^ have also been associated with SLID-resistance. However, no individual variant within these 18 genes was present in more than one AMK-R or CAP-R isolate, after removing variants present in at least one susceptible isolate (Table S5). Variants present in one or more susceptible isolates also did not significantly associate with resistance in this isolate set. There were however eleven unique mutations within the *whiB7* gene and its promoter, each carried by a separate unexplained KAN-R isolate, and no other isolates (Table S5). WhiB7 regulates *eis*^35^, which is in turn associated with KAN-R.

### WhiB7 Variants in Aggregate Associate with Kanamycin Resistance

Different variants in the same gene can cause the same change in phenotype^36^. If multiple rare variants in the same gene cause resistance, they may be missed by a genome wide association study on individual variants. To account for this, we measured the association between SLID-R and the variants in each *M. tuberculosis* gene in aggregate (Table 8). For each gene we identified all variants carried exclusively by resistant isolates, then counted the number of unexplained resistant isolates that carried at least one such variant in that gene (Table 8). No gene had more than 10 such isolates. To estimate how frequently mutations in the gene putatively cause unexplained resistance, this count was then divided by the total number of unexplained resistant isolates with any variant in that gene. For KAN, the *whiB7* promoter had the strongest signal, with seven unexplained KAN-R isolates carrying resistance-exclusive mutations (Table 8, Table S5). Each of the seven isolates carried a different *whiB7* promoter mutation, unique to that isolate. Meanwhile, five unexplained CAP-R isolates carried CAP-R exclusive mutations in the *thrB* promoter (Table 8). Mutations in the *thrB* promoter showed a similar signal for AMK-R, as did mutations in *ppe51* (Table 8). Beyond the top three genes and promoters, the proportion values for all three SLIDs dropped off noticeably.

**Table 8.**
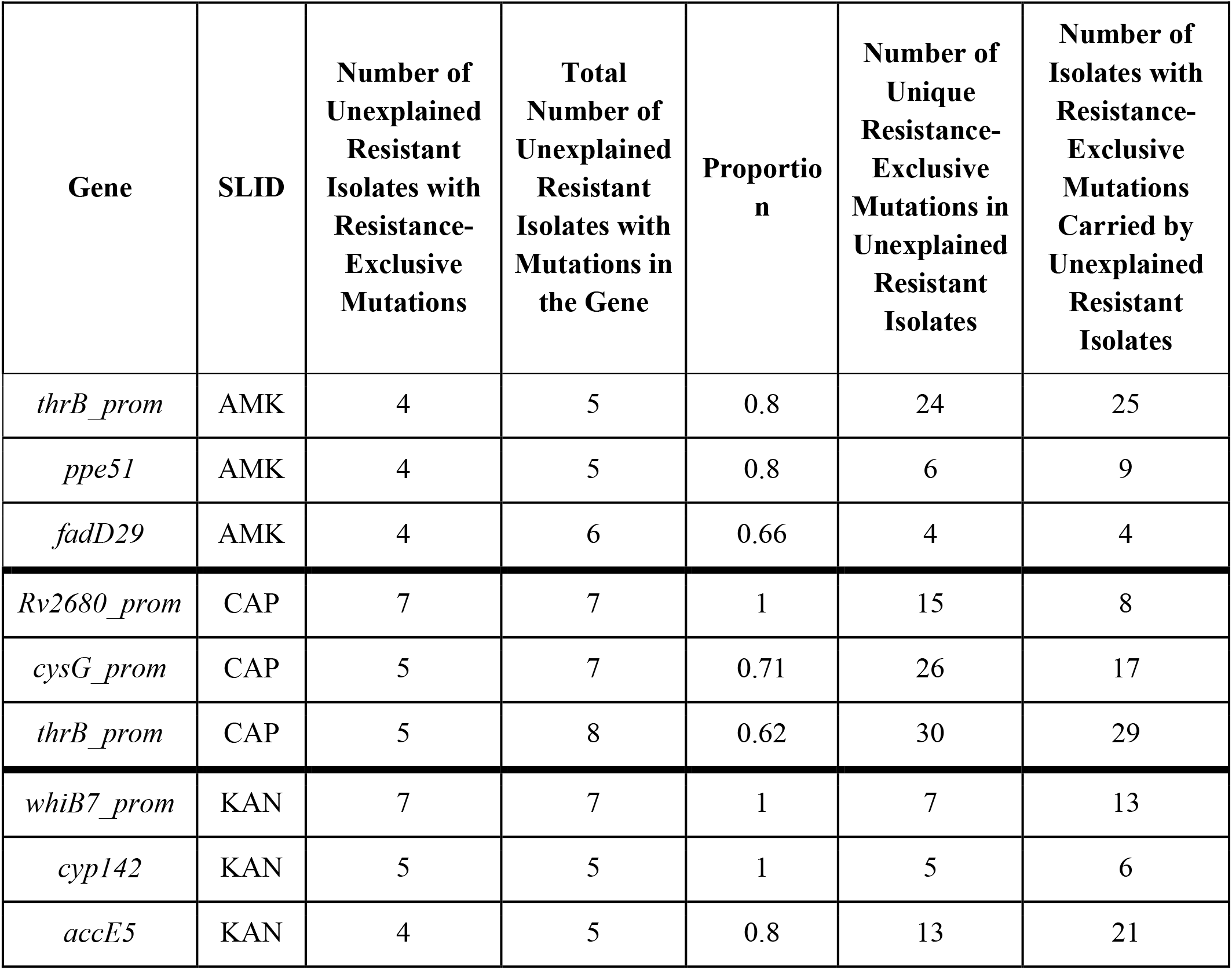
Gene-based genome-wide association study, top genes. “Number of Unexplained Resistant Isolates with Resistance-Exclusive Mutations” for each SLID and each gene counts the number of isolates that have unexplained resistance to that SLID and carry a mutation in that gene which is not carried by any isolate susceptible to that SLID. Genes below the mean for this count were removed (4 for AMK, 5 for CAP, and 3 for KAN). “Total Number of Unexplained Resistant Isolates with Mutations in the Gene” for each gene and each SLID counts the number of isolates that have unexplained resistance to that SLID and carry any mutation in that gene. “Proportion” is the first count divided by the second. Genes associated with first-line drug resistance were excluded.

| Gene | SLID | Number of Unexplained Resistant Isolates with Resistance-Exclusive Mutations | Total Number of Unexplained Resistant Isolates with Mutations in the Gene | Proportion | Number of Unique Resistance-Exclusive Mutations in Unexplained Resistant Isolates | Number of Isolates with Resistance-Exclusive Mutations Carried by Unexplained Resistant Isolates |
| --- | --- | --- | --- | --- | --- | --- |
| *thrB_prom* | AMK | 4 | 5 | 0.8 | 24 | 25 |
| *ppe51* | AMK | 4 | 5 | 0.8 | 6 | 9 |
| *fadD29* | AMK | 4 | 6 | 0.66 | 4 | 4 |
| *Rv2680_prom* | CAP | 7 | 7 | 1 | 15 | 8 |
| *cysG_prom* | CAP | 5 | 7 | 0.71 | 26 | 17 |
| *thrB_prom* | CAP | 5 | 8 | 0.62 | 30 | 29 |
| *whiB7_prom* | KAN | 7 | 7 | 1 | 7 | 13 |
| *cyp142* | KAN | 5 | 5 | 1 | 5 | 6 |
| *accE5* | KAN | 4 | 5 | 0.8 | 13 | 21 |

## Discussion

While known markers in *eis* promoter and *rrs* associated strongly with SLID-R, 111 SLID-R isolates lacked known markers in this study of 1184 clinical *M. tuberculosis* isolates. Some of the discordant isolates may be due to errors in phenotypic DST. Phenotypic/genotypic DST results sometimes disagree^37^ and have shown an error rate as high as 2.2% for AMK^38^. These discordant isolates could also suggest the existence of rare, alternative mechanisms of resistance. However, identifying rare mechanisms is difficult, as no single variant is carried by enough isolates for a strong association with resistance. To overcome this difficulty, we searched for rare mechanisms using a gene-based approach.

The known marker *rrs:*A1401G was ubiquitous and strongly associated with cross-resistance to all three SLIDs (Figure 2). However, while the variant was carried by 246 CAP-R isolates, it was also carried by 22 CAP-S isolates (Table 4). The discordant isolates may still have low level resistance to CAP, as a prior study reported a wide range of CAP MIC values (8 to 40 μg/ml) among clinical isolates carrying *rrs:*A1401G^39^. However, the cause of this variable MIC is still unknown. The same study reported that mutagenesis of reference strains consistently resulted in a 40 μg/ml CAP MIC, suggesting the inconsistency was due to the genetic background of the clinical isolates rather than the mechanism of *rrs:*A1401G itself^39^. However, we observed no mutations common to the genetic background of our 22 discordant isolates.

The known marker *rrs:*G1484T was rare, carried by only four clinical isolates (Table 4), all cross-resistant. The low prevalence of *rrs:*G1484T has been reported previously, despite a high MIC^14,40^. In a mutagenesis study on *M. smegmatis, rrs:*G1484T mutants grew slower than *rrs*:A1401G mutants, suggesting this rarity is due to a fitness cost of *rrs*:G1484T^41^. However, a later study found that *rrs*:G1484T conferred no growth disadvantage to the *M. tuberculosis* reference strain H37Rv, though it did confer a disadvantage to a strain of the Beijing F2 sublineage^39^. The full extent of the fitness cost of *rrs*:G1484T across clinical isolates is unknown.

While *eis* promoter mutations are generally known to cause KAN-R, mutagenesis experiments have shown that the *eis* promoter mutation *eis*:C-14T reduces susceptibility to AMK as well^15,42,43^. In 7H10 media, *eis*:C-14T mutants previously had MIC values (3 mg/L)^15^ between the current and previous WHO approved critical concentrations (2 mg/L and 4 mg/L, respectively)^44^. In our study DST were performed on MGIT at the unchanged critical concentration for that media (1 mg/L) and gave inconsistent results for isolates with *eis*:C-14T (11 AMK-R and 18 AMK-S, Table 3). These inconsistent results are likely due to the critical concentration’s proximity to the MIC of this variant. Though the genetic background of the different isolates could also be involved through epistatic effects.

KAN-R was most often explained by *rrs:*A1401G (Table 4), except among isolates from Moldova, which were enriched for known markers in the *eis* promoter (Table 5). Moldova is representative of countries in the region of the former Soviet Union, where this geographic trend has been reported previously^16^. The prevalence of *eis* promoter mutations in the region is thought to be the result of extensive use of KAN^16^.

While known markers in *eis* promoter and *rrs* associated strongly with resistance, no other mutations within these genes significantly associated with SLID-R in this isolate set (Figure 3). The mutation *rrs*:C1402T was infrequent and carried by both resistant and susceptible isolates (Figure 3). Mutagenesis has previously shown that *rrs*:C1402T reduces susceptibility to a level near the critical concentration^39^. Both AMK-R and AMK-S isolates carried *rrs*:C1402T in a recent WHO study^43^, and the marker is considered resistant by the GenoType MTBDRsl platform (Hain Lifescience).

The streptomycin resistance marker *rrs:*A514C was enriched among isolates with the known SLID-R marker *rrs:*A1401G (OR = 6.0, p = 2.424e-11, Fisher’s Exact Test). Streptomycin has previously been widely used in treatment regimens when first-line antibiotics fail^2^. It is thus likely that the association between these two variants is due to past use of streptomycin on MDR-TB strains that were later treated with SLIDs.

The known *eis* promoter and *rrs* mutations (Table 1) were not carried by 21 of the 270 KAN-R isolates (Table 3), leaving the genetic basis of their resistance unexplained. Of these, seven isolates carried *whiB7* promoter mutations (Table 8, S5), though these mutations were unique in each isolate. Thus, while no single *whiB7* promoter mutation associated strongly with KAN-R, their aggregate signal suggests *whiB7* promoter mutations are an alternative mechanism of KAN-R. This finding in clinical isolates is supported by prior mutagenesis experiments. Increased expression of *whiB7* causes low level streptomycin and KAN-R in H37Rv mutants^33^, while deletion of *whiB7* in *Mycobacterium abscessus* lowers MIC to erythromycin, tetracycline, streptomycin, and AMK^46^. The *whiB7* gene encodes a transcriptional activator that regulates ribosome protection and efflux pump genes^46^. Moreover, *whiB7* regulates *eis*, providing it a plausible mechanism for KAN-R^33^.

Similarly, mutations in the gene *ppe51* collectively associated with AMK-R (Table 8). PPE51 mediates membrane transport and loss of function mutations in *ppe51* have recently been shown to cause resistance to the drug candidate 3,3-bis-di(methylsulfonyl)propionamide^25^ and *ppe51* knockout increased susceptibility to pyrazinamide^47^. Mutations in *ppe51* have also been observed in an experimentally evolved CAP-R strain, though this strain also carried a *tlyA* frameshift mutation^48^. The associated *ppe51* mutations in our data were not found in isolates with *tlyA* mutations, except for the ubiquitous synonymous mutation *tlyA*:L11L. The presence of *ppe51* mutations here in AMK-R isolates, in the absence of canonical markers or *tlyA* mutations, supports their potential role in resistance. Mutations in the *thrB* promoter also collectively associated with both AMK-R and CAP-R (Table 8). However, unlike *whiB7* and *ppe51, thrB* promoter mutations lack a plausible mechanism of resistance. ThrB is a homoserine kinase involved in threonine biosynthesis and is essential for virulence and *in vitro* growth^49^, but there is no known connection between this pathway and SLID-resistance.

The known SLID-R markers are accurate predictors of resistance. However, they still do not explain all SLID-R cases. Rare, alternative mechanisms, such as *whiB7* promoter mutations, are likely responsible for these unexplained SLID-R isolates. For molecular diagnostics to fully replace phenotypic diagnostics, these rare mechanisms must also be understood. Finding these rare mechanisms will require sequences from larger sets of unexplained resistant isolates, and more sensitive methods of association, such as the machine learning approaches employed previously^21^ or the gene-based aggregate method employed here. This method independently corroborated the association between *whiB7* promoter mutations and KAN-R^46^, and identified a new association between AMK-R and mutations in *ppe51*, a gene previously implicated in resistance to other compounds^25,47,48^.

## Materials and Methods

### Sample Collection

As part of a previous study, 323 clinical *M. tuberculosis* isolates were collected for long read PacBio sequencing^18^. There were 89 isolates that originated from Hinduja National Hospital (PDHNH) in Mumbai, India, 89 that came from the Phthisiopneumology Institute (PPI) in Chisinau, Moldova, 48 which were from the Tropical Disease Foundation (TDF) in Manila, Philippines, and 97 that were from the National Health Laboratory Service of South Africa (NHLS) in Johannesburg, South Africa. All raw sequences were uploaded to NCBI’s sequence read archive (SRA) database under the Bioproject accession PRJNA353873. An additional 10 *M. tuberculosis* clinical isolates were collected from the Supranational Reference Laboratories in Stockholm and Antwerp. These isolates were originally genotyped with a Hain Lifescience Genotype MTBDRsl line probe assay^9^, and were chosen for sequencing due to discordance between their genotype and phenotypic DST for any SLID. Another 851 whole genome sequences were downloaded from NCBI’s SRA database using SRA Toolkit’s fastqdump^50^. These 851 downloaded raw reads were previously sequenced on Illumina short read platforms^51–54^.

### Phenotypic Drug-susceptibility Testing

DST for the PacBio sequenced isolates was performed on the BACTEC mycobacterial growth indicator tube (MGIT) 960 platform (BD Diagnostic Systems, Franklin Lakes, NJ, USA) using the 2008 WHO recommended critical concentration of 1.0 mg/L (AMK) and 2.5 mg/L (CAP/KAN) as described in previous studies^18,55,56^. DST for Illumina sequenced isolates were also tested on MGIT 960 using contemporary WHO recommended critical concentrations, as described previously^51–54^. As of 2018, the recommended critical concentrations for AMK, CAP, and KAN remains 1.0 mg/L, 2.5 mg/L, and 2.5 mg/L, respectively^44^. Bacterial isolates were excluded from analysis if DST data was not available for at least one SLID.

### DNA Extraction and Sequencing

The DNA of all 333 isolates collected for long-read PacBio sequencing, including those from the WHO Supranational Reference Laboratories in Stockholm and Antwerp, and from NCBI’s SRA database were extracted as described in a previous study^57^. The SMRT sequencing protocol was described previously^58,59^. 64 isolates were later re-sequenced due to low coverage. DNA extraction for the 851 downloaded public genomes was previously described^51–54^. The downloaded genomes were sequenced on Illumina Genome Analyzer, MiSeq, or HiSeq platforms.

### Genome Assembly, Alignment and Variant Calling

Genome assembly, alignment, and variant calling methods are described in Supplementary Information. Briefly, PBHoover^60^ aligned 64 SMRT sequenced isolates to H37Rv and called variants. Later, 269 SMRT sequenced isolates were de novo assembled with canu^61^ or HGAP2 (https://github.com/PacificBiosciences/Bioinformatics-Training/wiki/HGAP-2.0) then their assembled genomes were aligned to reference strain H37Rv using dnadiff (v1.3)^62^ for variant calling, with the output converted to VCF format by a custom script, mummer-snps2vcf (https://gitlab.com/LPCDRP/mummer-extras/-/blob/master/src/mummer-snps2vcf). Reads from Illumina sequenced isolates were aligned to H37Rv using bowtie2 (v2.2.4)^63^, then variants were called with VarScan2 (v2.3)^64^.

### Lineage Identification

MIRU-VNTR and spoligotyping were previously performed on the initial 323 SMRT sequenced isolates collected^55^. The ten isolates sent from Stockholm and Antwerp, and the 851 downloaded Illumina sequenced isolates, underwent MIRU-VNTR and spoligotyping with MiruHero, a custom Python script (https://gitlab.com/LPCDRP/miru-hero). MiruHero used the rule based criteria from TB-Insight^65^ to classify lineages.

### Identifying Known Resistance Conferring Mutations

After variant calling, known SLID-R markers were searched for in the VCF file of each clinical isolate. The *eis* promoter mutations C-14T, C-12T, and G-10A are known KAN-R markers, and the *rrs* mutations G1484T and A1401G are known resistance markers to all three SLIDs^18^. The genomic positions and orientation of *rrs* and the *eis* promoter were noted in Table S6^66^. Known markers and phenotypic DST were used to estimate the sensitivity and specificity of predicting resistance to each SLID using known markers. A 95% confidence interval was calculated for sensitivity and specificity estimates using the score method with continuity correction^28^.

### Genome Wide Association

For each of the three drugs, separate genome wide association studies were performed using a custom Python script (https://gitlab.com/LPCDRP/gwa) to identify novel markers for alternative mechanisms of resistance. To remove the overriding signal of known resistance markers, our GWAS excluded isolates that had their resistance explained by known markers (Table 1). This exclusion was necessary to avoid confounding associations with potential alternative mechanisms of resistance. Isolates with additional *eis* promoter resistance markers (Table 1) were excluded in our KAN-R analysis for similar reasons. We calculated the sensitivity and specificity of each variant’s prediction of resistance to each SLID (the proportion of unexplained resistant isolates with the variant, and proportion of susceptible isolates without the variant, respectively). For each SLID we identified variants absent from susceptible isolates and ranked them by the number of unexplained resistant isolates that carried them. The same protocol was performed using a subset of genes previously implicated in SLID-R.

In the gene-based association, for each SLID and each gene we counted the number of unexplained resistant isolates with at least one variant in that gene that was absent from isolates susceptible to that SLID. Genes below the mean for this count were removed. This count was then divided by the total number of unexplained resistant isolates with any variant in that gene. Genes with known resistance markers to first line drugs were excluded, as most SLID-R isolates are also resistant to first-line drugs due to prior treatment with first-line drug regimens.

## Supplementary Information Availability

Supplementary Tables, Figures, and methods are available at https://doi.org/10.5281/zenodo.5720106.

## Acknowledgements

Phenotypic confirmation of ten isolates without common molecular mechanisms of resistance was performed by Mr. Solomon Ghebremichael in the Department of Microbiology at the Public Health agency of Sweden in Stockholm.

